# Bridging local and global dynamics: a biologically grounded model for cooperative and competitive interactions in the brain

**DOI:** 10.1101/2025.07.09.663817

**Authors:** Borja Mercadal, Maria Guasch-Morgades, Lucia Mencarelli, Giacomo Koch, Giulio Ruffini, Alzheimer’s Disease Neuroimaging Initiative

## Abstract

Functional brain networks exhibit both cooperative and competitive interactions, yet existing models—assuming purely excitatory long-range coupling—fail to account for the widespread anti-correlations observed in fMRI. Starting from a laminar neural mass framework, where each mass comprises distinct slow (alpha-band) and fast (gamma-band) oscillatory pyramidal subpopulations (P1 and P2), we show how laminar-specific long-range excitatory projections across neural mass parcels can give rise to both cooperation and competition via cross-frequency envelope coupling. We demonstrate that homologous connections across parcels (e.g., P1→P1 or P2→P2) induce positive correlations between the infra-slow amplitude fluctuations of alpha band envelopes in each parcel, as well as in the simulated fMRI BOLD signals. Conversely, heterologous connections (P1→P2) induce negative correlations. We tested this mechanism by building personalized whole-brain models for a cohort of 60 subjects in two steps. First, we inferred signed inter-parcel generative effective connectivity directly from resting-state fMRI using regularized maximum-entropy (Ising) models. Then we connected laminar neural masses to simulate BOLD dynamics by implementing positive and negative Ising connections via homologous and heterologous projections, respectively. Ising-derived cooperative/competitive connectivity modeling faithfully reproduced both static and dynamic functional connectivity patterns, as well as gamma power-BOLD correlation and partial alpha power-BOLD anticorrelation–outperforming structurally constrained and cooperative-only variants. This further demonstrates that functional data alone suffices to infer individualized connectivity. Together, these results provide a biologically grounded mechanistic model on how long-range excitatory circuits and local cross-frequency interactions shape the balance of cooperation and competition in large-scale brain dynamics.

**Highlights:**

- We introduce a biologically grounded mechanism for brain-wide cooperation and competition, based on laminar-specific cross-frequency coupling (CFC) between alpha and gamma oscillations, mediated solely by excitatory long-range projections targeting different cortical layers.
- Our models can reproduce key empirical observations, including the negative correlation between alpha power and BOLD, the positive correlation between gamma power and BOLD, and the presence of local laminar cross-frequency interactions consistent with invasive and EEG findings.
- Models generate BOLD signals autonomously, without recourse to external stochastic inputs.
- We introduce a method for personalized Ising modeling from Ising data using a sparsity (L1) constraint to infer signed connectivity from limited BOLD data.
- We validate the proposed mechanism in a cohort of 60 subjects using subject-specific generative whole-brain models, which not only improve the replication of static functional connectivity but also accurately capture dynamic spatiotemporal brain state transitions.
- Our whole-brain models display biologically realistic local dynamics, with laminar neural mass models preserving plausible EEG-like alpha and gamma oscillations while aligning with large-scale BOLD patterns—offering a unifying framework that bridges microscale laminar physiology and macroscale functional connectivity.

## 1 Introduction

The ability of the brain to dynamically engage and disengage distributed networks is fundamental to the orchestration of perception, cognition, and behavior. Both at rest and during task performance, brain regions exhibit temporally structured patterns of coordination, giving rise to networks that transiently synchronize and desynchronize in response to evolving cognitive demands.^1–3^ These patterns of inter-regional interactions, commonly measured using functional magnetic resonance imaging (fMRI) as functional connectivity (FC), are characterized not only by positive (cooperative) correlations, reflecting coordinated fluctuations in activity, but also by negative (competitive) correlations, where increased activity in one region is associated with decreased activity in another.^1, 4–7^

While both cooperative and competitive interactions are well-documented features of largescale brain dynamics, the neural mechanisms that generate these interactions remain poorly understood. Prevailing models of large-scale brain dynamics have predominantly focused on cooperative interactions,^8–12^ which are typically attributed to long-range excitatory anatomical connections that facilitate the synchronization of activity across distant cortical regions. However, this framework does not adequately explain the robust and reproducible anti-correlations observed in empirical FC data, which suggest the presence of competitive processes that counteract global cooperation. Recent computational modeling studies have incorporated competitive (negative) interactions to better replicate the complex topology of empirical FC.^13–15^ However, such modeling approaches lack a clear physiological grounding, treating competition as a mathematical abstraction rather than as an emergent property of brain circuits.

A growing body of evidence points to the interplay of fast and slow brain rhythms as a key mechanism for shaping interactions within and between brain regions. In particular, cross-frequency coupling (i.e. the dynamic interaction between slow, alpha, and fast, gamma, oscillations) has been proposed as a mechanism for coordinating hierarchical information flow across cortical circuits.^16–20^ According to predictive coding theories, alpha oscillations mediate top-down predictions, while gamma oscillations reflect bottom-up prediction errors, with their interaction supporting the comparison of expectations and sensory inputs.^18, 19, 21^ Importantly, these oscillations are layer-specific, with alpha and gamma rhythms generated in different cortical layers and associated with distinct feedforward and feedback projections.^19, 22–24^

These findings suggest that laminar-specific long-range projections interacting with local oscillatory circuits may be crucial in dynamically balancing cooperation and competition across the brain. However, how these mechanisms scale to shape large-scale network dynamics remains unclear. In particular, while it is known that long-range cortical connections target specific layers,^25, 26^ and that alpha and gamma oscillations reflect distinct functional states,^27, 28^ it is not well understood how these anatomical and physiological features give rise to the dynamic functional architecture observed in fMRI, including widespread anti-correlations.

Intriguingly, although data fitting of functional connectivity data using generative effective modeling using Ising^29, 30^ or Hopf^13^ models reveals the need for both positive and negative connections across brain parcels, a biologically realistic circuital implementation of these interactions has not yet been provided. To address this gap, we propose a biologically grounded mechanistic framework that links laminar-specific long-range excitatory projections and cross-frequency interactions to the emergence of both cooperative and competitive interactions in the FC landscape. Building on anatomical studies of layer-specific projections and theoretical work on predictive coding,^18, 20^ we hypothesize that long-range excitatory projections modulate alpha and gamma oscillations across regions, generating either cooperative (correlated) or competitive (anticorrelated) dynamics mediated by cross-frequency coupling^18, 20, 31, 32^ depending on their laminar targeting. This, in turn, can yield the full range of positive and negative correlations observed in large-scale brain networks.

We adopt a multi-scale modeling approach to investigate the neural mechanisms underlying these interactions in brain networks. We begin by the single cortical column, modeled with a laminar neural mass model (LaNMM),^31, 32^ to explore how the interplay between fast (gamma) and slow (alpha) oscillatory dynamics shapes neural activity and the corresponding simulated BOLD signals. By extending this analysis to pairs of interconnected columns, we demonstrate how layer-specific long-range excitatory projections can give rise to either positive or negative correlations between regions, depending on their laminar targeting. Building on these insights, we then construct subject-specific whole-brain generative models with both cooperative and competitive interactions to test whether the local mechanisms identified at the columnar level can reproduce empirical FC patterns. Finally, we analyze the spatiotemporal dynamics generated by these whole-brain models, evaluating their ability to reproduce the brain’s dynamic functional repertoire. Through this progression from local circuit dynamics to large-scale brain networks, our work provides a biologically grounded framework for understanding how cooperation and competition emerge from cortical microcircuits and shape macroscopic brain activity.

## 2 Methods

### 2.1 Empirical data

#### 2.1.1 Data acquisition

In the present study, neuroimaging data from a total of 60 subjects with no known neurological condition was used. Among those 60 subjects, the data of 23 was obtained from our ongoing clinical trial (NCT06826261).^33^ The study has been approved by the review board and ethics committee of the Santa Lucia Foundation. All patients or their relatives or legal representatives have provided written informed consent. Imaging was conducted in the Memory Clinic of the Santa Lucia Foundation (Rome, Italy) with a 3T Siemens Magnetom Prisma scanner with a 64-channel head-neck coil. T1-weighted MRI images were acquired using standardized MPRAGE protocols. Key acquisition parameters included TR = 2500 ms, TE = 1.67 ms, voxel size = 1×1×1 mm, and flip angle = 8°. Diffusion weighted MRI (dMRI) images were acquired using an Echo Planar Imaging sequence (EPI) (TR=3400 ms, TE=80 ms, flip angle (FA) = 90°, matrix size=116×116, slice thickness=1.8 mm). dMRI mages were obtained with a total of 48 diffusion sampling directions, the b-value was 1000 s/mm2. Finally, resting-state fMRI images were acquired using standard echo-planar blood oxygenation level-dependent (BOLD) imaging.

Key parameters included TR = 980 ms, TE = 30 ms, flip angle =63°, voxel size = 2.8×2.8×2.8 mm, and scan duration ∼ 8 minutes.

The data from the remaining 37 subjects was obtained from the Alzheimer’s Disease Neuroimaging Initiative (ADNI) database (adni.loni.usc.edu). The ADNI was launched in 2003 as a public-private partnership, led by Principal Investigator Michael W. Weiner, MD. The original goal of ADNI was to test whether serial magnetic resonance imaging (MRI), positron emission tomography (PET), other biological markers, and clinical and neuropsychological assessment can be combined to measure the progression of mild cognitive impairment (MCI) and early Alzheimer’s disease (AD). Detailed imaging protocols and scanner-specific adjustments are available in the ADNI3 documentation.

#### 2.1.2 Data processing

The T1-weighted MRI images were processed using the recon-all pipeline of Freesurfer 6.0 http://surfer.nmr.mgh.harvard.edu/.^34^ This pipeline was used to annotate the voxels in the cortex^35^ following the Desikan-Killiany parcellation atlas^36^ in addition to the voxels belonging to 8 sub-cortical areas (Thalamus, caudate, putamen, pallidum, hippocampus, amygdala, nucleus accumbens-a and the cerebellum). This resulted in a total of 84 parcels (34 cortical parcels per hemisphere and 8 sub-cortical parcels per hemisphere).

To process the fMRI BOLD images, first, fMRIPrep^37^ was used to co-register with the structural MRI images, apply slice timing correction and estimate the head motion artifacts. Then, ANTS^38^ togheter with Freesurfer^34^ were used to register the images to MNI space and map the parcellation previously obtained from the structural MRI into the fMRI images. Then, the average activity per parcel was calculated by averaging the activity in the voxels within each parcel. Finally, global signal regression (GSR) was applied by regressing out the average signal intensity across all brain parcels.

Diffusion MRI data was processed to calculate the subject’s structural connectome (SC). Preprocessing steps included denoising, artifact correction (e.g., eddy currents), and estimating the fiber orientation density using constrained spherical deconvolution. Probabilistic tractography was then performed with MRtrix3^39^ generating 20 million streamlines constrained to start and terminate in gray matter parcels defined by the MRI parcellation. ANTS^38^ togheter with Freesurfer^34^ were again used to register the images to MNI space and map the parcellation previously obtained from the structural MRI. Finally, the SC matrix was obtain by quantifying the number of streamlines connecting each pair of parcels.

#### 2.2 Laminar Neural Mass Model

In the present study, the local oscillatory dynamics of each brain region were simulated using the LaNMM. This model was was originally introduced to model the laminar generation of oscillations in two bands.^31, 32^ It has been used recently to model fast interneuron dysfunction in Alzheimer’s Disease,^40^ the impact of pshychedelics in cortical dynamics,^41^ and predictive coding.^20^ The rich dynamical landscape of the model has been recently been provided.^42^ The model combines two well-established neural mass models, the Jansen-Rit (JR)^43^ and a modified Pyramidal Interneuron Gamma (PING) model,^44^ to simulate slow and fast oscillatory dynamics within a cortical column (see Figure 1a). The JR sub-model generates alpha-band (8–12 Hz) oscillations and operates at ∼10 Hz under baseline conditions, while the modified PING sub-model produces gamma-band (30–70 Hz) oscillations with a baseline frequency of ∼40 Hz. The reciprocal coupling of these models displays marked cross-frequency interactions, including signal to envelope coupling (SEC) and envelope to envelope (EEC, also known as amplitude-amplitude coupling) coupling of slow and fast oscillations,^20^ aligning with observed experimental data where alpha rhythms influence gamma activity.^18, 24^ In particular, the architecture yields a positive SEC and a negative EEC in the operating regime derived from macaque data.^20, 32^

**Figure 1.**
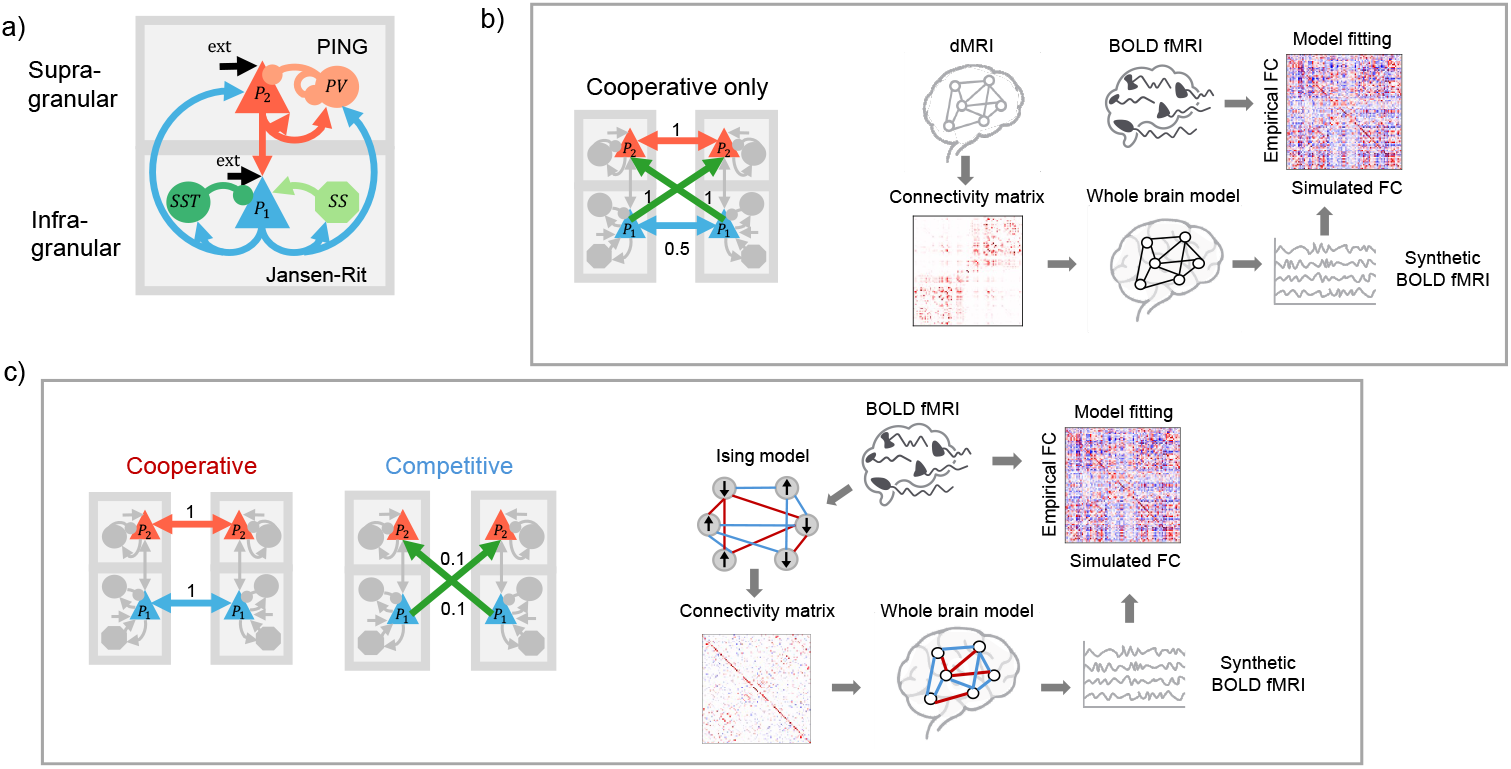
a) Schematic of the neural populations and the connections between them in the LaNMM. Inhibitory synapses are represented with a ball while excitatory synapses are represented with an arrow. b) Diagram of the pipeline used to create subject-specific whole brain models from structural data and using cooperative interactions only. The connections between populations and the relative weights used in the long range connections are depicted in the left. c) Diagram of the pipeline used to create subject-specific whole brain models with cooperative and competitive interactions. The connections between populations and their relative weights for cooperative and competitive interactions are shown in the left. Note that generative connectivity matrices in this case were obtained from BOLD fMRI data using Ising models^30^

The JR model consists of a population of Pyramidal neurons (*P*_1_), a population of excitatory cells (e.g., Spiny stellate cells *SS*), and a population of slow GABAergic interneurons (e.g., Somatostatin-expressing cells *SST*, such as Martinotti cells). The PING model consists of two populations, a Pyramidal (*P*_2_) one and a fast GABAergic interneuron one (e.g., Parvalbumin-positive cells *PV*, such as basket cells). The equations and parameters of the model can be found in section S1 of the Supplementary materials.

The JR populations are associated with infra-granular neural populations. In particular, the pyramidal population *P*_1_ can be linked to Layer 5 pyramidal cells (L5PC). In contrast, the populations in the PING are associated with supra-granular populations. Specifically, the pyramidal population *P*_2_ can be linked to Layer 2-3 pyramidal cells (L2-3PC). These associations between neural populations in the model and cortical layers were derived through electrophysiological data assimilation^32^ and are also consistent with experimental findings.^22–24, 28^

### 2.3 Generative effective connectivity with Ising models

#### 2.3.1 Spin-glass model

Subject-specific effective connectivity matrices were generated following an approach based on the Ising spin-glass model from Sherrington and Kirkpatrick^45^ that has already been used by several studies in neuroscience.^14, 30, 46, 47^ Ising models treat each brain region as a unit that can be ‘active’ or ‘inactive’, and can be used to infer the coupling strength between each pair of regions to reproduce observed co-activations in neuroimaging data.

The spin-glass model is defined by the energy or Hamiltonian of the lattice of *N* spins with pairs of parcels *i* and *j* (*i, j ϵ* [1, *N*], *i*≠*j*), given by

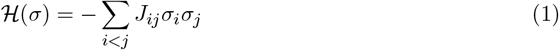

where *σ*_*i*_ denotes the orientation of each spin in the lattice (±1) and *J*_*ij*_ is the coupling matrix or Ising connectivity. The sum over *i < j* denotes a sum over all pairs of spins (with pairs counted once). We assume that self-connections are zero (*J*_*ii*_ = 0) and that *J* is symmetric (*J*_*ij*_ = *J*_*ji*_).

In the context of neuroscience, elements such as neurons, columns or brain regions can be modeled by spins (i.e., with two states, up or down, on or off) with pair interactions. Specifically, in the present study *σ*_*i*_ represents the binarized state of each brain parcel as active (+1) or inactive (−1). Thus, Ising models can be derived from BOLD fMRI data by transforming the time series in each parcel into a binary format. In the present study, for each parcel, the median of the time series (after global signal regression) was used as a threshold for binarization. That is, BOLD time series samples were assigned a value of +1 if its value was greater than the threshold and −1 otherwise.

Following the Maximum Entropy Principle,^48^ spin models aim to identify the least-biased probability distribution that reproduces observed constraints, such as pairwise spin correlations derived from data. The coupling matrix *J* can be estimated using an approximation to maximum likelihood as described in by Ezaki et al.^49^ Briefly, *J* is derived by maximizing the pseudo-likelihood function,

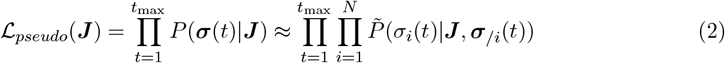

where 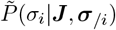 is the modeled probability distribution for a specific spin given the states of all the others defined as

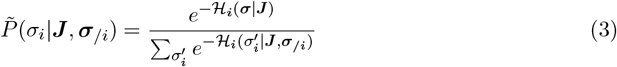

with ℋ_*i*_(***σ***|***J***) from Equation 1.

Using this approximation, the pseudo-likelihood can be maximized by gradient ascent

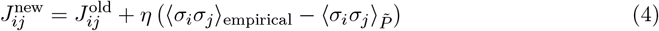

where 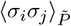 is the two-point correlation function with respect to distribution 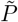,^49^

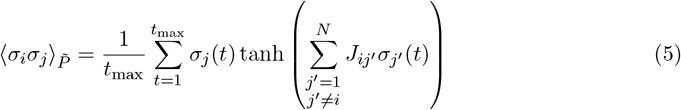

#### 2.3.2 Structural connectivity constrained connectomes

The maximization problem in Ising models for deriving brain connectivity can be degenerate, especially considering the limited number of time points per parcel available in BOLD fMRI data. A possible approach to handle such degeneracy is to use the structural connectivity derived from dMRI data as a constraint in the maximization problem.^46^ Briefly, the approach is based in adding the squared difference between the Ising connectivity *J* and the sctructural connectome *SC* as a penalty in the optimization scheme by defining the constrained log-pseudolikelihood function as

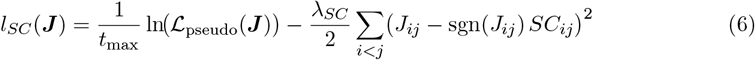

Again, a gradient ascent procedure can be used to maximize *l*_*SC*_(***J***) which yields the following updating scheme,

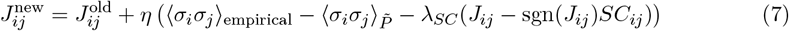

The value of the penalty *λ*_*SC*_ was adjusted to achieve a mean correlation of 0.95 between the Ising-derived and the empirical FC matrices (see section S2 in the Supplementary materials).

#### 2.3.3 Sparsely constrained connectomes

An alternative approach to the use of structural data is to constrain the optimization of *J* using an L1 norm,^50^ favoring sparse solutions,

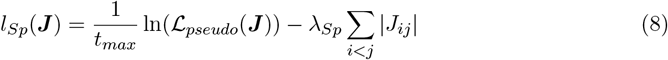

In this case the gradient ascent procedure to maximize *l*_*SC*_(***J***) yields the updating scheme

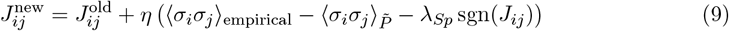

As in the previous section, the value of the penalty *λ*_*Sp*_ was adjusted to have a mean correlation of 0.95 between the FC matrix derived from the Ising model and the FC matrix derived from the empirical fMRI BOLD data (see section S2 in the Supplementary materials).

### 2.4 Generation of synthetic BOLD fMRI data

The BOLD time series of a brain region were calculated using a well established modeling framework that combines the so-called Balloon model^51^ and a hemodynamic model^52^ that relates the neural activity with changes in the blood flow and blood volume. All model equations and parameters are listed in section S3 of the Supplementary materials.

Despite its widespread use, there is not a well established way of computing the “neural activity” that drives this model from the variables of generative whole brain models. Local ATP consumption for ion-pump activity and neurotransmitter cycling drives the neurovascular response underlying the BOLD signal.^53–55^ In addition, it has been shown that synaptic inputs rather than spiking outputs are the main driver for such metabolic demand.^53, 55, 56^ From an energetic perspective, the metabolic demand associated with post-synaptic currents scales with two main factors:

1. The synaptic activity, governed by the pre-synaptic firing rates, *φ*_*j*_, and the number of connections, *C*_*i*←*j*_.
2. The post-synaptic potentials amplitude and kinetics which depend on the synaptic gains *A*_*i*←*j*_ and rate constants *a*_*i*←*j*_ associated with each synapse.

Thus, from our models we can construct a metabolic drive, *m*(*t*), by summing over all neural populations *i* in a parcel and over all pre-synaptic populations *j*:

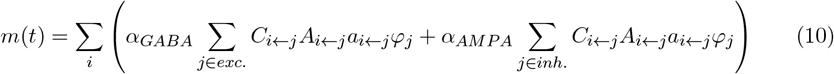

where the subscripts *i* ← *j* indicate a synapse from the neural population *j* to the neural population *i* and the parameters *α*_*GABA*_ and *α*_*AMPA*_ convert synaptic currents (Gabaergic and Ampaergic respectively) to ATP consumption. Previous studies have used different values to weight the contribution of inhibitory and excitatory synapses^57, 58^ or even further refined approaches.^59^ For simplicity, we only considered the contribution of the excitatory synapses (*α*_*GABA*_ = 1 and *α*_*AMPA*_ = 0) which allows further simplifying 10 since all excitatory synapses in our models have the same synaptic gain and rate constant values. Thus, the absolute neural activity *X*(*t*) in a parcel was simply calculated as:

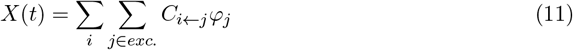

In the supplementary materials section S4 we show that our results are robust to this choice.

Finally, the relative neural activity, *x*(*t*), was calculated by normalizing *X*(*t*) to its mean value over time 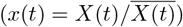. Then, *x*(*t*) was used as an input to the Balloon model described in section S3 of the Supplementary materials.

### 2.5 Subject-specific generative whole brain models

To investigate the role of cooperative and competitive long-range interactions in whole-brain dynamics, we constructed two types of subject-specific whole-brain models based on their long range connections: i) models with cooperative interactions only using connectivity strengths from the SC derived from the subject’s dMRI data, and ii) models accommodating for both cooperative and competitive interactions, with the connectivity type and strengths between regions derived from the Ising effective connectivity models described in 2.3.

The pipelines used to personalize these two types of whole-brain models, as well as the specific architecture of long-range connections, are illustrated in Figure 1 b and and c.

To build the generative whole brain models, a LaNMM^32^ was placed in each brain region (parcel), and these LaNMM units were connected via long-range connections. The pattern of long-range connections was designed to reflect plausible cortico-cortical communication schemes derived from predictive processing theories^18, 19, 60^ and supported by anatomical and physiological evidence. Specifically, experimental data indicate that L5PC form long-range excitatory projections to both infra- and supra-granular layers of distant cortical columns (feedback projections), while L23PCs primarily target granular layers, with subsequent relay to L23PCs (feedforward projections).^25, 26^ Since granular populations are not explicitly modeled in the LaNMM, the L23PC to L23PC pathway was implemented as a simplified representation of the translaminar feedforward connections mediated via the grnaular layer.

Based on the above, three variants of long-range connections between parcels *i* and *j* were implemented:

1. 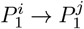: infra-granular to infra-granular (feedback) connection.
2. 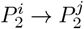: supra-granular to supra-granular (feedforward) connection.
3. 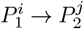: infra-granular to supra-granular (feedback) connection.

For simplicity, this connectivity scheme is symmetric and only employs long-range excitatory coupling. Excitatory pyramidal (glutamatergic) neurons account for roughly 80–85% of cortical cells and send extensive axons across cortical and subcortical targets, forming the backbone of inter-areal communication.^61^ Long-range excitatory projections in neocortex arise almost exclusively from pyramidal neurons and overwhelmingly target other pyramidal cells rather than interneurons. Quantitative mapping in somatosensory cortex shows that L2/3 and L5 pyramidal neurons send axons whose boutons predominantly innervate the dendrites of target-area pyramidal neurons, with only a minor fraction contacting inhibitory cells.^62, 63^ Within each cortical area, intratelencephalic and pyramidal-tract pyramidal cells form reciprocal pyramidal-to-pyramidal loops at densities up to an order of magnitude higher than projections onto interneurons. In contrast, long-range GABAergic neurons represent only a small minority of cortical projection cells (*<* 5%) and mostly modulate distant circuits via disinhibition rather than direct inhibition of principal cells.^64^ Focusing on pyramidal-to-pyramidal pathways therefore captures the dominant mode of cortico-cortical communication and simplifies our model by accounting for the bulk of long-range excitatory interactions.

The connectivity strength, *C*_*j,l*←*i,k*_ of each long-range connection from population *k* in parcel *I* and population *l* in parcel *j* is modeled as

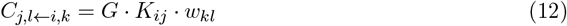

where *K*_*ij*_ is the connectivity strength from the connectivity matrix (in absolute value), *w*_*kl*_ is the relative weight of the connection between populations and the *G* is a global coupling factor that scales the overall connectivity strength.

In the models with cooperative and competitive interactions, the signs of the connectivity values in the connectomes were used to select between the two connectivity schemes in Figure 1c (positive values associated with cooperative and negative values associated with competitive).

These specific laminar connections were designed to either induce a correlation or an anticorrelation between the fast and slow oscillators, and consequently modulate BOLD fMRI correlations between regions, as demonstrated in the Results section 3.1. Namely, *P*_1_ → *P*_1_ and *P*_2_ → *P*_2_ connections are meant to induce a correlation in the amplitude envelope oscillations with the goal of yielding a positive correlation between BOLD signals (cooperative interaction). In contrast, *P*_1_ → *P*_2_ connections are meant to anti-correlate the alpha and gamma envelopes, and consequently yield an anti-correlation in the BOLD signals (competitive interaction).

The relative weights of the three connection types were optimized (see Supplementary Material Section S5) to maximize the Pearson correlation coefficient (PCC) between empirical and simulated BOLD FC matrices.

Constant external inputs were set to the *P*_1_ and *P*_2_ populations (no stochastic noise was added to the model). In order to preserve the LaNMM-specific local oscillatory dynamics observed in a single-column LaNMM, the values of these were adjusted for each value of *G*. The adjustment was made to ensure that the mean input received by each LaNMM unit remained consistent with the operating regime of an isolated column. This adjustment compensates for the changes induced by the net incoming long-range excitatory input to each pyramidal population in the laminar model.

The global coupling parameter *G*, which scales the overall strength of long-range interactions between brain regions, was adjusted individually using grid search for each subject-specific whole-brain model to optimize the match between simulated and empirical FC. For each candidate value of *G*, the full system of coupled LaNMM units was simulated for 600 seconds, with a sampling rate of 500 Hz. The differential equations of the model were integrated using a fourth-order Runge-Kutta (RK4) method. During each simulation, the firing rates of excitatory populations in each parcel were recorded and subsequently used to generate synthetic BOLD signals via the hemodynamic Balloon-Windkessel model, as described in Section 2.4. GSR was applied to the generated BOLD signals, and they were subsequently used to compute the synthetic FC matrix, defined as the PCC between the time series of all pairs of brain regions. The empirical FC matrix for each subject was derived analogously from the processed resting-state fMRI data. Finally, the value of *G* that maximized the PCC between the synthetic and empirical FC matrices was selected as the optimal coupling strength for that subject.

### 2.6 Leading Eigenvector Dynamics Analysis

The model parameters for the different approaches were adjusted to match the empirical static FC matrix. To further assess how well the spatiotemporal dynamics are captured by these models, the Leading Eigenvector Dynamics Analysis (LEiDA) framework was employed to analyze the resulting BOLD time series. This approach was first introduced by Cabral et al.^65^ and has later been exploited for data assimilation in the framework of whole brain modeling.^66, 67^ Deco et al.^66^ introduced the Probabilistic Metastable Substates framework, which provides a probabilistic description of the spatiotemporal dynamics and, after identifying the brain states of interest, the resting state FC can be assimilated to a walk on a discrete Markov chain.

The Python toolbox PyLeida was used to cluster the leading eigenvectors from the empirical data (after GSR) with various cluster numbers between two and ten. The optimal number of clusters was selected based on various metrics such as the silhouette score, the Dunn score, and the Davies-Bouldin score. In addition, the correlation between the FC states and the seven cerebral intrinsic functional networks estimated by Yeo et al.^2^ was analyzed following the same approach as in Vohryzek et al.^68^ This was done to ensure that the selected cluster is physiologically meaningful.

To assess how well the different models reproduced empirical spatiotemporal dynamics, a similar approach to that of Deco et al. (2019)^66^ was followed. Once the optimal number of clusters was found, both the empirical and synthetic data were partitioned into the FC states. To do so, the cosine distance between each data point and the cluster centroids was calculated, and the nearest cluster label was assigned to the data point. Following partitioning, the probabilities of each FC state, *π*_*i*_, were calculated as the fractional occupancy (FO) of the states (for every subject and model or empirical data). That is, the number of samples (i.e., time points) assigned to the FC state divided by the total number of samples.

The symmetrized Kullback–Leibler (KL) distance between the empirical, 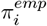, and synthetic, 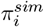, FC state probabilities was used to assess how well the different modeling approaches reproduce the FO of the different states:

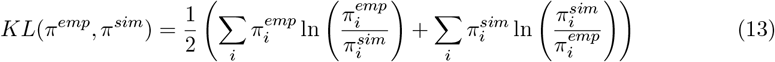

with *i* being the FC state index.

For each subject (and for each group of interest—empirical data or one of our modeling approaches), we also estimated a *k* × *k* transition-probability matrix

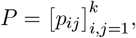

where *p*_*ij*_ = Pr{*X*_*t*+1_ = *j* | *X*_*t*_ = *i*}. Matrices were computed exactly as in Vohryzek *et al*. (2020).^68^ Under the usual assumption that *P* is ergodic (irreducible and aperiodic), there exists a unique stationary distribution

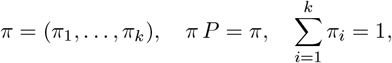

which characterizes the long-run occupancy of each state.^69^ We then computed the *entropy rate* by

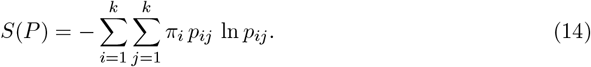

To assess how well the models capture the transition patterns, the absolute difference of the synthetic and empirical entropy rates was calculated as:

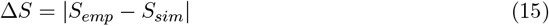

Large differences imply that either the models are too random or too predictable compared to empirical data, while low values mean that transition dynamics are correctly reproduced by the models.

## 3 Results

### 3.1 LaNMM and BOLD signals

We first analyzed the intrinsic dynamics of a single LaNMM and their relationship with the BOLD signals generated by the column. Figure 2a shows a sample of the membrane potentials of the pyramidal populations in the model and the amplitude envelopes in the alpha band (for *P*_1_) and the gamma band (for *P*_2_). The figure also shows the infra-slow fluctuations in the amplitude envelopes in the alpha and gamma bands *E*_*α*_ and *E*_*γ*_, together with the BOLD signal generated by the column. Here, we will use ‘infra-slow fluctuations’ (ISFs) to refer to *<* 0.1 Hz modulations of the amplitude envelope of band-limited neural oscillations.^70^ The membrane potentials of *P*_1_ and *P*_2_ display a cross-frequency coupling: 1) the envelope of *P*_2_ correlates with the phase of *P*_1_, and 2) the ISF *E*_*α*_ and *E*_*γ*_ show a negative correlation (EEC). In addition, BOLD signal correlates with *E*_*γ*_ and is anti-correlated with *E*_*α*_. This BOLD signal correlation with the frequency bands is consistent with the literature that establishes a correlation with the gamma envelope^53, 71, 72^ and an anti-correlation with the alpha envelope.^72–74^

**Figure 2.**
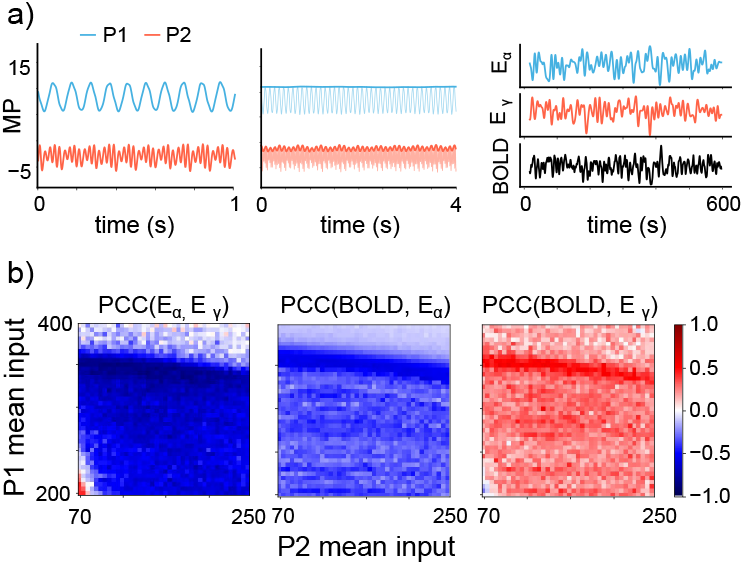
Single Laminar Neural Mass Model. a) LaNMM dynamics: the left plot displays the membrane potentials of the pyramidal populations in a single LaNMM. The center plot shows the amplitude envelopes of the membrane potentials computed using the norm of the Hilbert transform after band pass filtering in the alpha band (*P*_1_) or the gamma band (*P*_2_). The right plot shows the infras-low fluctuations in the amplitude envelopes *E*_*α*_ and *E*_*γ*_ together with the BOLD signal generated by the column. *E*_*α*_ and *E*_*γ*_ were computed by low pass filtering (cutoff frequency of 0.1 Hz) the amplitude envelopes in the alpha and gamma bands. b) Pairwise correlations between the infra-slow fluctuations in the alpha, *E*_*α*_, and in the gamma band, *E*_*γ*_, as well as with the BOLD time series as a function of the mean external inputs to *P*_1_ and *P*_2_ populations.

Remarkably, the cross-frequency EEC (negative) as well as the BOLD correlation/anti-correlation with the ISFs of the amplitude envelopes *E*_*α*_ and *E*_*γ*_ are robust over a wide range of external inputs to the *P*_1_ and *P*_2_ populations as shown in Figure 2b.

Next, we investigated how long-range excitatory projections between two LaNMM columns shape inter-parcel BOLD correlations. We tested different bi-directional connections, targeting either homologous (*P*_1_ → *P*_1_ or *P*_2_ → *P*_2_) or heterologous (*P*_1_ → *P*_2_) pyramidal populations, and examined the resulting correlations between BOLD signals. The results are displayed in Figure 3 together with the inter-parcel correlation between the ISF *E*_*α*_ and *E*_*γ*_ of the columns.

**Figure 3.**
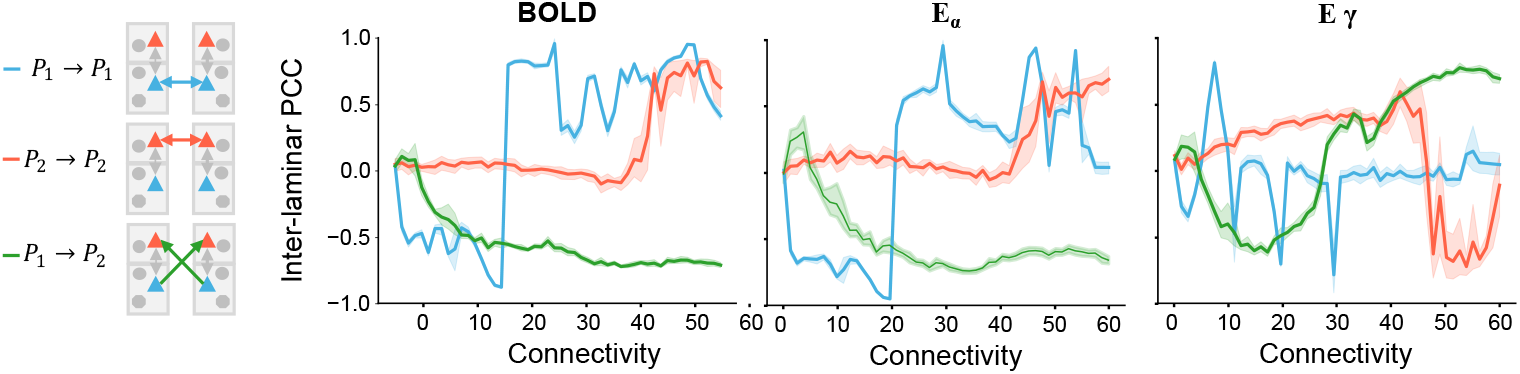
Connected Laminar Neural Mass Models. Inter-laminar Pearson Correlation Coefficient (PCC) of the BOLD signals and the amplitude envelopes *E*_*α*_ and *E*_*γ*_ of two LaNMM columns bi-directionally connected with different schemes (*P*_1_ → *P*_1_, *P*_2_ → *P*_2_ and *P*_1_ → *P*_2_). The results displayed are the average of 10 runs with different random seeds and the shaded areas show the standard deviation.

The results show that homologous connections induce correlations between the ISF in the alpha-band, leading to positive BOLD correlations. Conversely, heterologous connections induce anticorrelated ISF in the alpha envelopes, generating negative BOLD correlations between columns.

Interestingly, although some of the BOLD correlations/anti-correlations are mediated by the gamma oscillators, the ISF of the gamma envelopes themselves do not consistently correlate or anti-correlate.

Altogether, these results show that the LaNMM architecture enables the cooperation and competition between cortical layers through intra-laminar cross-frequency coupling. BOLD signals correlate if *E*_*α*_ across columns are correlated. This correlation can appear via a direct connection between the alpha oscilators (*P*_1_ → *P*_1_), but also mediated by the EEC when connecting the gamma oscillators of the two columns (*P*_2_ → *P*_2_). Similarly, BOLD signals anti-correlate when there is an inter-laminar anti-correlation in *E*_*α*_. This can be achieved by connecting the alpha oscillator of one column to the gamma oscillator of the other (*P*_1_ → *P*_2_). In that scenario, *E*_*α*_ in one column modulates *E*_*γ*_ in the other, which then modulates (negatively) the *E*_*α*_ of the column yielding an inter-laminar negative correlation of the *E*_*α*_. Remarkably, these findings demonstrate that both cooperative and competitive interactions between brain regions (i.e., positive or negative correlations in BOLD) can emerge from long-range excitatory projections alone.

### 3.2 Generative effective connectivity

Prior to build whole brain models, we generated subject-specific connectivity matrices with three different approaches: 1) the structural connectome (SC) derived from the dMRI data, 2) Ising derived connectomes constrained with the structural connectivity (Ising-SC) and 3) Ising derived connectomes with a sparsity constraint (Ising-L1). It is important to note that, unlike the SC, Ising connectomes feature both positive (i.e., cooperative) and negative (i.e., competitive) connections.

Figure 4 outlines the differences between the connectivity matrices obtained with these approaches. A guide is provided to help the reader interpret the connectivity matrices in Figure 4a. Figure 4b displays the connectomes obtained with these three approaches for an exemplary subject. Two main traits can be noticed: first, the ising connectomes display high connectivity between homotopic regions (i.e., mirror areas of the two brain hemispheres), which is not present in the SC and, second, inter-hemispheric connectivity is much sparser in the SC than in the ising connectomes.

**Figure 4.**
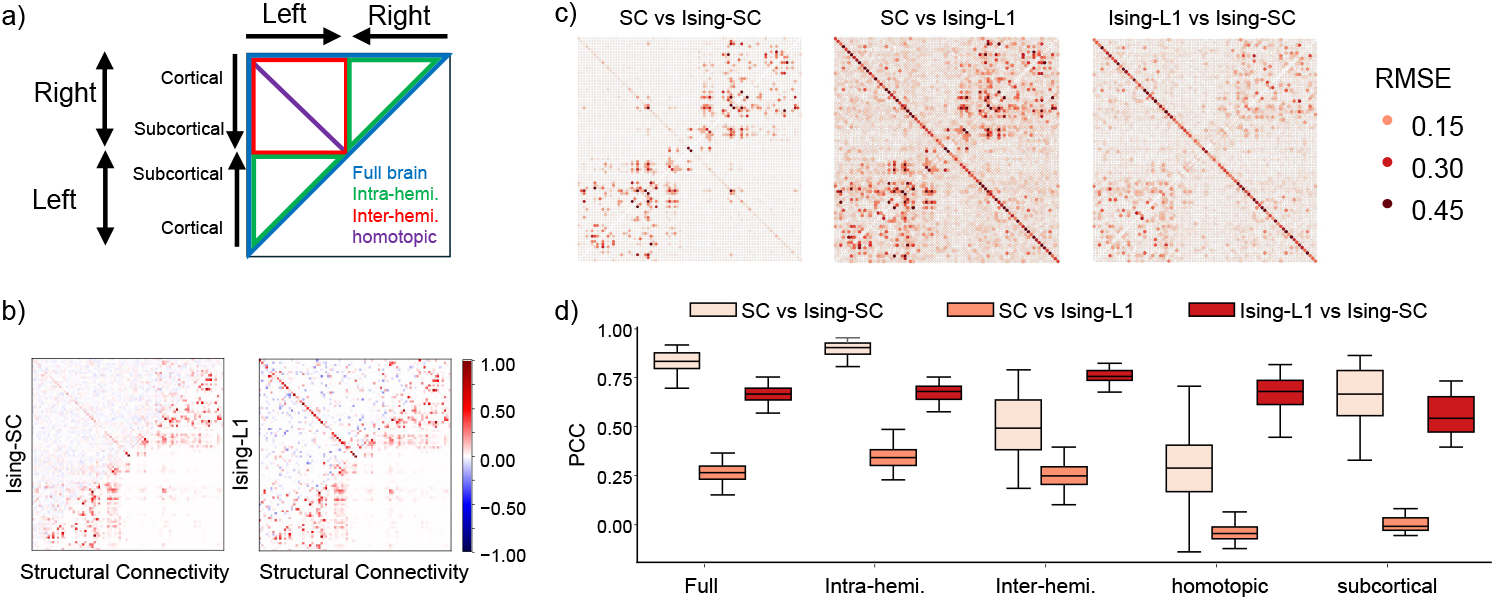
Generative effective connectivity (GEC) matrices. a) Guide to interpret connectivity plots: Full connectivity matrix (in blue) can be dissected into intra-hemispheric (green triangles) and inter-hemispheric(red square). With the arrangement used in this study, the line *y* = *x* corresponds to the matrix diagonal, and the cross-diagonal corresponds to the homotopic connections. Note that the order of the parcels is inverted between hemispheres. b) Connectivity matrices obtained with the two Ising approaches (upper triangles) with the structural connectivity (lower triangles) for an exemplary subject. c) Average root mean squared error (RMSE) between connectivity matrices across all subjects and for each element in the connectivity matrices. d) Pearson’s correlation coefficient (PCC) between connectivity matrices across all subjects. The box plots show the PCC over the full matrix and over various relevant parts.

To provide an overview of the main differences between connectomes, we computed the root mean squared error (RMSE) between the matrices for each pair of parcels across all subjects (see Figure 4c). In addition, we computed the PCC between the matrices per subject for the full matrix and also for various relevant subsets of connections (see Figure 4d).

The greatest discrepancies between SC and Ising-SC occur in intra-hemispheric edges: the SC graph is sparser but has stronger connections compared to Ising-SC. We also observe pronounced differences in subcortical links and homotopic pairs. Overall, SC and Ising-SC remain highly correlated, largely due to their intra-hemispheric agreement, while correlations drop for inter-hemispheric and homotopic connections. This reflects the SC’s inherently sparse crosshemisphere wiring and weak homotopic strength, which the Ising-SC model augments with additional coupling.

Among all pairwise comparisons, Ising-L1 vs. SC shows the largest discrepancies in both RMSE and Pearson correlation. These differences are most pronounced in the SC’s dense connectivity clusters and in homotopic and inter-hemispheric links. Moreover, the PCC analysis reveals substantial mismatches in subcortical connections as well.

Finally, the comparison between Ising-L1 and Ising-SC connectomes shows a high overall correlation, indicating that these two generative models produced largely similar results. The main differences occur in inter-hemispheric connections and can be seen clearly in homotopic regions, where Ising-L1 exhibits stronger connectivity. However, despite the large RMSE, we obtained a high PCC between the connectomes in the inter-hemispheric regions. Connections involving subcortical parcels emerge as the subset with the lowest correlation, while the rest of the connectome displays high correspondence between the two models. The RMSE analysis also highlights differences in certain intra-hemispheric connections, particularly in clusters associated with high connectivity in SC, likely due to the impact of the structural connectivity constraint in the Ising-SC model.

Across all comparisons, both Ising variants augment the sparse inter-hemispheric and homotopic edges of the empirical SC, leading to the largest RMSE and lowest Pearson correlations in those cross-hemisphere and subcortical links. Intra-hemispheric connections remain largely preserved, yielding high overall correlations with SC. Between the two models, Ising-L1 and Ising-SC align closely overall—their main divergence being that Ising-L1, unrestrained by structural weights, produces even stronger homotopic and inter-hemispheric couplings than Ising-SC.

### 3.3 Generative whole brain models

Next, we integrated the subject-specific connectivity matrices into whole-brain models to test how well they reproduce empirical FC. This was done by placing a LaNMM in each brain parcel and connecting them using the three connectome types described above. Models were optimized to maximize the PCC between synthetic and empirical FC matrices. Crucially, in the case of Ising connectomes, the signs of the matrix entries determined the selection of cooperative (positive) and competitive (negative) long-range interactions.

Figure 5a shows the synthetic FC of the best fits obtained with the three connectivity matrices against the empirical FC for an exemplary subject. Figure 5b summarizes the overall fitting results for the three families of connectivity matrices. The models with cooperative and competitive interactions yield substantially better fits with the empirical data than the models with cooperative interactions only. Interestingly, although the differences between the two Ising connectomes are not as large, they display a statistically significant difference. The connectomes obtained with the Ising and a sparsity constraint yield better fits than those obtained with the structural connectivity as a constraint. It is important to note that both Ising frameworks were adjusted to achieve similar goodness-of-fit to the empirical FC at the Ising model level. Thus, constraining the Ising model to be similar to the structural connectome did not have a positive impact on the resulting whole-brain models.

**Figure 5.**
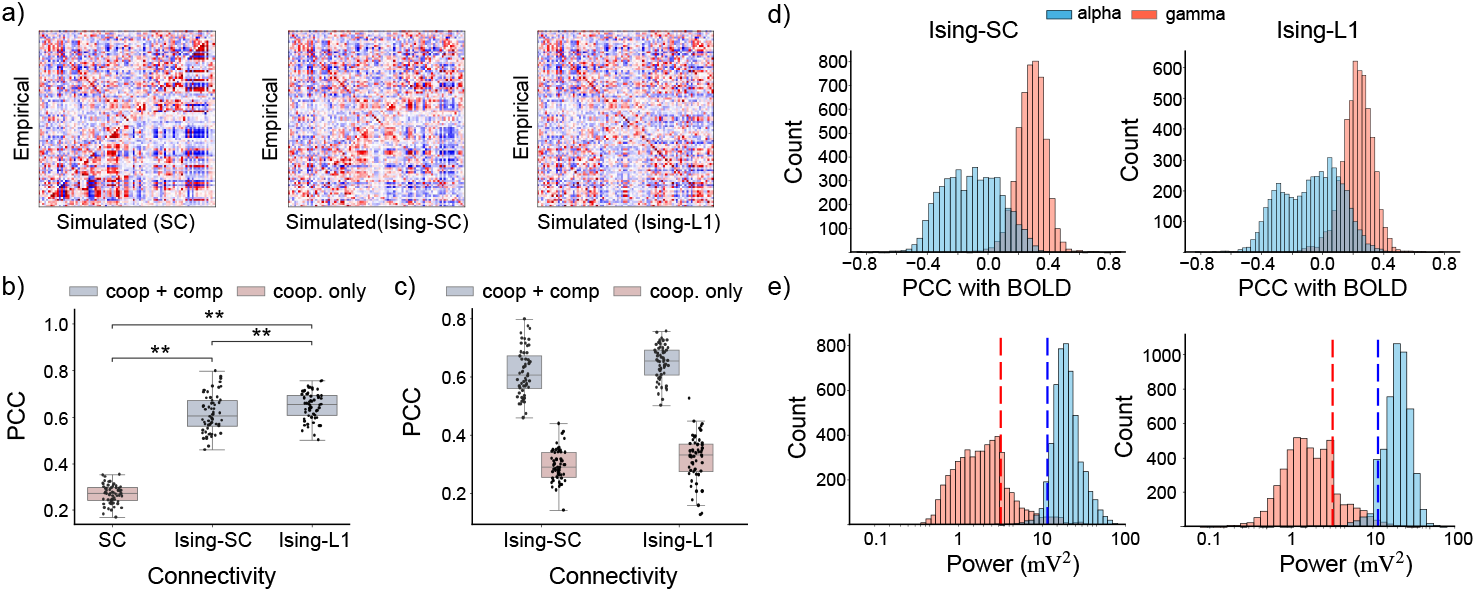
Subject specific whole brain models. a) Comparison between the empirical (upper triangle) and synthetic (lower triangle) FC matrices of a sample subject for the models using each of the three connectivity matrices. Note that while the model using the structural connectivity only had cooperative interactions, the models using the Ising connectomes had both cooperative and competitive couplings. b) Pearson’s correlation coefficient between the empirical and the synthetic FC matrices in each individual subject for the three types of connectivity matrices. c) Model fits obtained using the connectomes when only cooperative interactions are used or when both types of interactions are included. d) Local (i.e. parcel-wise) correlation between the BOLD signals and the ISF in the amplitude envelopes *E*_*α*_ and *E*_*γ*_ in the Ising-SC (left) and the Ising-L1 (right) models. e) Local power of the membrane potential in the alpha band (for *P*_1_) and in the gamma band (for *P*_2_). Vertical lines show the values of a single column LaNMM.

To rule out the possibility that such strikingly better fits obtained with the models using cooperative and competitive interactions are solely due to the (unsigned) connectivity strength, we repeated the process with the Ising connectomes but only using cooperative interactions using the absolute value of the connectomes. Namely, we modeled all long-range connections using the connectivity scheme and the relative weights from Figure 1b. The results are displayed in Figure 5c. Even though the results of the cooperative only models improve compared to those obtained with the SC, the models with both types of interactions still yield significantly better fits.

Finally, it is worth noting that these generative models not only reproduce the whole brain dynamics (i.e., the FC), but also have realistic local oscillatory dynamics as well as BOLD correlation/anti-correlation patterns with the ISF of the amplitude envelopes. Figure 5d shows the correlations between BOLD and the ISF *E*_*α*_ and *E*_*γ*_ in all parcels for the two models using cooperative and competitive interactions. Aligned with the results shown for the single column LaNMM, BOLD signals largely correlate with *E*_*γ*_ and, to a lesser extent but preferentially, anticorrelate with *E*_*α*_. In addition, the mean powers of the membrane potentials of the pyramidal populations in the alpha and gamma bands are similar to those obtained with a single LaNMM column as shown in Figure 5e.

### 3.4 Spatiotemporal dynamics assessment

To assess the ability of the different whole-brain models to reproduce the empirical spatiotemporal dynamics of brain activity, we applied the Leading Eigenvector Dynamics Analysis (LEiDA) framework^11^ to both empirical and synthetic BOLD fMRI data.

First, we determined the optimal number of FC states by evaluating clustering solutions ranging from 2 to 10 clusters using multiple clustering quality metrics, including the Dunn score, Silhouette score, Davies-Bouldin score, and distortion score (Figure 6a). These scores collectively indicated that three clusters provided the most robust and consistent partitioning of the data. In addition, we selected this number of clusters as it produced FC states that showed high correspondence with well-established functional networks, as reported by Yeo et al.^2^

**Figure 6.**
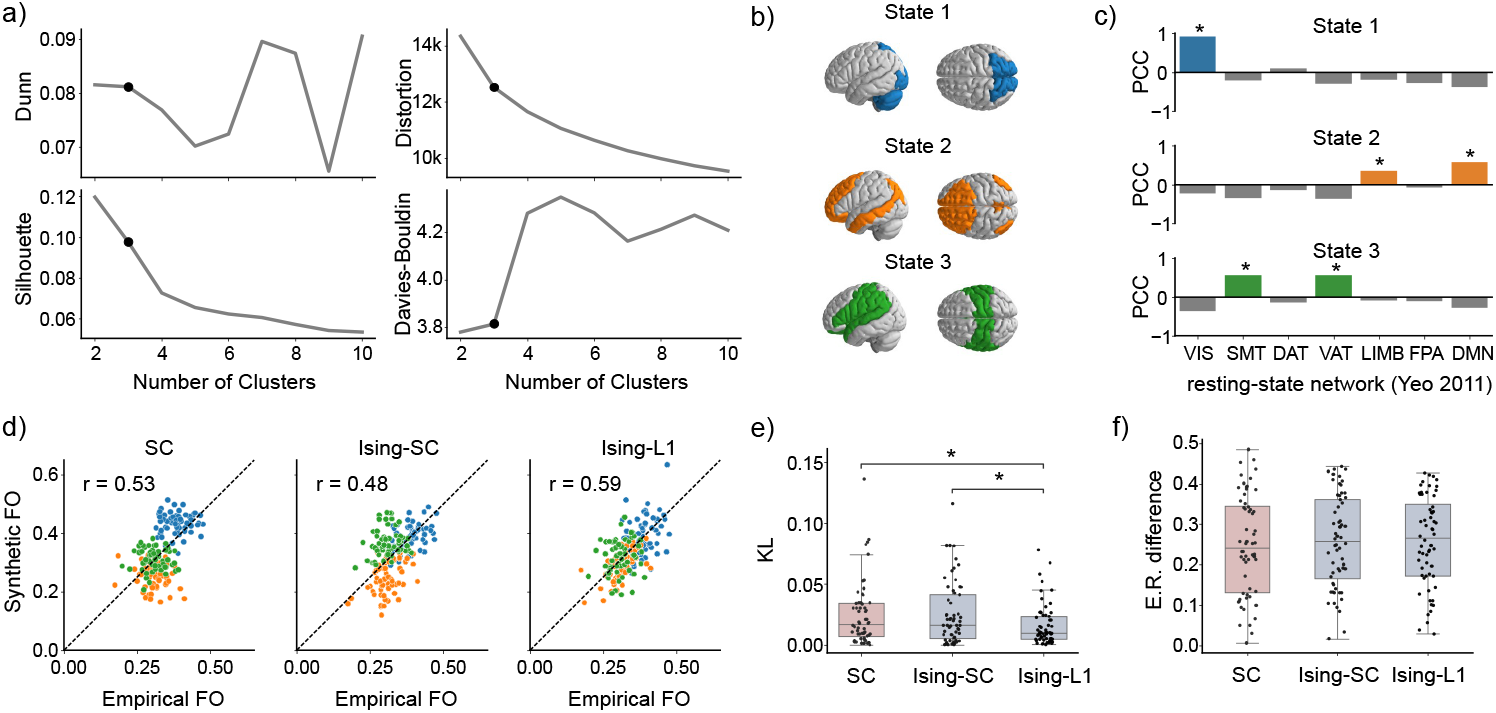
Spatiotemporal dynamics assessment. a) Clustering performance metrics obtained when applying the LEIDA algorithm. b) Representation of the centroids of the FC states of the optimal clustering on the cortical surface. c) Overlap between the FC states of the selected clustering and the seven functional resting state networks from Yeo et al.^2^ computed as the Pearson’s correlated coefficient (VIS: Visual, SMT: Somatomotor, DAT: Dorsal attention, VAT: Ventral attention, LIMB: Limbic, FPA: Frontoparietal, DMN: Default mode). Asterisks indicate statistical significance (*p <* 0.01). d) Scatter plots of the empirical and the synthetic fractional occupancies for each subject, FC state and modeling approaches. Each point represents a subject and the colors represent the different FC states: blue for State 1, orange for State 2, and green for State 3. The dashed line has a slope of 1 and an intercept equal to zero. e) Kullback–Leibler distance between synthetic and empirical data for the different groups. Asterisks (*p <* 0.05, Wilcoxon signed-rank test). f) Entropy rate difference between synthetic and empirical data for the different groups (no statistically significant differences found between groups using a Wilcoxon signed-rank test)

Furthermore, the analysis of the spatial patterns of the FC state centroids revealed meaningful overlap with canonical resting-state networks (Figure 6b, c). Specifically, State 1 exhibited a high correlation with the Visual network (*r* = 0.92, *p <* 0.01), State 2 showed significant correlations with both the Default Mode Network (*r* = 0.59, *p <* 0.01) and the Limbic network (*r* = 0.36, *p <* 0.01)), and State 3 was correlated with both the Somatomotor (*r* = 0.56, *p <* 0.01) and Ventral Attention networks (*r* = 0.56, *p <* 0.01).

Next, we compared the fractional occupancy (FO) or percentage of time the brain spends in each recurring FC state, between empirical and simulated data (Figure 6d). The models incorporating both cooperative and competitive interactions, particularly those using Ising-L1 connectomes, showed the highest agreement with empirical FO distributions, as indicated by the global PCC between simulated and empirical FOs. In contrast, models using only structural connectomes and cooperative interactions tended to deviate more substantially from the empirical distribution.

These observations are quantitatively confirmed by the symmetrized Kullback–Leibler (KL) divergence between the empirical and synthetic FOs (Figure 6e). The Ising-Sp models exhibit significantly lower KL distances than both the cooperative-only models and the structurally constrained Ising-SC connectomes (*p <* 0.05, Wilcoxon signed-rank test), indicating that Ising-Sp best reproduces the probabilistic structure of the brain dynamics. In contrast, entropy-rate differences –which reflect the overall randomness of transitions– do not differ significantly between groups (Figure 6f). Thus, the lower KL divergence shows that Ising-Sp captures the occupancy of states more faithfully. In contrast, the similar entropy rates imply that all models approximate the overall frequency and unpredictability of state transitions to a comparable degree. Likely reflecting that all models are more predictable than experimental data to the same extent.

It is worth noting that the advantage of incorporating competitive interactions remained consistent across a range of cluster numbers (3 to 7), as shown in the Supplementary Materials (Section S6). Specifically, Ising-L1 models showed significantly lower KL divergence compared to SC, regardless of the number of clusters. However, the difference between the Ising-SC and the Ising-L1 models is attenuated for large cluster numbers. The results regarding entropy rate differences were instead not robust to the number of clusters selected. Surprisingly, the SC-derived model shows significantly smaller entropy rate differences compared to the other groups for a larger number of clusters (5 to 7).

Overall, the model combining both cooperative and competitive interactions (Ising-L1) most faithfully reproduces empirical spatiotemporal dynamics. Although entropy-rate remains similar across models (reflecting equivalent global randomness in the transition dynamics), only Ising-L1 matches the state probabilities. In contrast, enforcing structural priors (Ising-SC) does not improve, and in fact slightly impairs, the ability of the generative models to replicate the empirical data.

## 4 Discussion

Our findings reinforce recent work^13–15, 29, 30^ demonstrating that competitive interactions are essential to reproduce key features of whole-brain dynamics, including anti-correlations and spatiotemporal variability. However, while previous modeling studies introduced competitive interactions as mathematical adjustments to improve model fits, our study goes further by providing a biologically grounded mechanism for such interactions, based on the interplay of fast and slow oscillatory circuits within cortical columns (i.e., the cross-frequency coupling). Using the LaNMM, we show that long-range excitatory projections targeting specific cortical layers can induce either positive or negative correlations between the ISF of the amplitude envelopes or BOLD signals in brain regions, depending on the location of synaptic targeting. In particular, we show that connections between homologous oscillatory circuits across parcels yield positive correlations while connections between heterologous oscillatory circuits yield negative correlations. This provides a mechanistic explanation, grounded in known anatomical projections and cross-frequency coupling, for the emergence of both cooperative and competitive interactions in large-scale brain networks relying only on long-range excitatory connections.

A central feature of our model is the EEC between infra-granular (alpha) and supra-granular (gamma) populations, whereby fluctuations in alpha power directly modulate the amplitude envelope of gamma oscillations. This negative EEC between *E*_*α*_ and *E*_*γ*_—high alpha envelope coinciding with low gamma envelope—recapitulates invasive recordings showing that alpha activity anti-correlates with gamma-band power in sensory cortex.^16–18, 27^

Furthermore, because the BOLD signal predominantly reflects metabolic processes tied to synaptic and spiking activity—which are most strongly driven by gamma-band oscillations— our mechanistic EEC link naturally yields the empirically observed negative correlation between alpha-band power and BOLD^72–76^ alongside the positive correlation between gammaband power and BOLD.^71–76, 76^ Thus, EEC in the LaNMM provides a biophysically grounded explanation for why alpha and gamma envelopes drive opposing BOLD fluctuations. In addition, this negative EEC plays a key role in the emergence of competitive interactions as it allows, via connections between heterologous oscillators, inducing anti-correlated ISF of the amplitude envelopes.

Remarkably, unlike single column neural mass models,^77^ the whole brain models can generate BOLD-like activity and similar fits to the FC empirical data even if no stochastic driving noise is added to the neural mass models (see supplementary materials section S7). Thus, our models are deterministic and although the results depend on the initial conditions the model fits are robust to them as shown in the supplementary materials.

In our models, long-range connections primarily modulate alpha-band envelopes, yielding cooperative and competitive interactions that shape inter-parcel BOLD correlations. In contrast, while gamma amplitude envelopes correlate with BOLD activity locally, they do not appear to be directly modulated by long-range interactions, even if the connections are mediated by gamma oscillators (i.e., via *P*_2_). This aligns with human MEG and EEG-fMRI studies showing that resting-state network interactions are predominantly mediated by slow oscillatory fluctuations, particularly in the alpha and beta bands, which display correlated activity across distant cortical areas.^78, 79^ In addition, animal models have shown that low-frequency oscillations and not gamma predominantly contribute to BOLD correlations between brain regions.^80^ In contrast, gamma-band power is tightly linked to local neuronal processing and metabolic demand, showing strong correlations with regional BOLD fluctuations but little evidence of inter-regional synchronization at rest.^71, 72, 76^

The proposed mechanism of excitatory projections producing competitive interactions is also consistent with anatomical evidence regarding the layer-specific targeting of long-range connections. Studies have shown that feedback and feedforward pathways target distinct cortical layers, with L5PCs projecting to both infra- and supra-granular layers in distant columns.^18, 25, 26^ How-ever, the specific populations mediating these effects are not fully established, and it remains unclear whether inhibitory populations might also be involved in long-range interactions, contributing to competition between regions.^81, 82^ Therefore, while our model proposes a plausible excitatory-only mechanism, alternative explanations involving disynaptic inhibitory pathways cannot be excluded.

A key strength of the modeling approach used in this study is the balance between realistic local and global dynamics. Our generative whole-brain models not only reproduce realistic large-scale dynamics (i.e., the static FC), but also maintain biologically plausible local oscillatory dynamics. This contrasts with prior studies that have introduced competitive or negative connections in whole-brain models to better match FC but lacked biologically interpretable mechanisms for such interactions or did not feature realistic local dynamics.^13–15^

In addition, to assess the validity of our whole-brain models beyond static FC, we employed LEiDA to evaluate the ability of the models to reproduce the spatiotemporal structure of BOLD activity. LEiDA has proven to be a powerful tool for characterizing dynamic FC and identifying recurrent FC states, with demonstrated sensitivity to clinical and experimental conditions.^65, 83, 84^ Our results show that models incorporating both cooperative and competitive interactions, particularly those using Ising connectomes with sparsity constraints (Ising-L1), best capture the distribution of brain states. This suggests that the inclusion of competitive dynamics is critical for reproducing the temporal richness of brain activity, consistent with prior work highlighting the importance of metastable transitions in brain dynamics.^67, 85^

We used Ising models to infer subject-specific generative effective connectivity matrices independently of the dynamical simulations used to model brain activity. This contrasts with other studies in which effective connectivity is adjusted within the same generative model that is fitted to functional data.^13, 15, 86, 87^ By separating the estimation of effective connectivity from the whole-brain simulation, our framework prevents potential circularity or model-specific biases and allows a clearer interpretation of competitive and cooperative interactions as reflecting intrinsic properties of the data rather than artifacts of model fitting. Moreover, the Ising approach offers flexibility in the use of structural connectivity: while it can incorporate anatomical priors when available, it also allows effective connectivity to be inferred solely from functional data and simplicity biases.

In fact, although this was not the main focus of the study, the finding that personalized wholebrain models can be generated independently of structural connectivity data emerges as an important and relevant outcome. While SC has traditionally been used to constrain whole-brain models, our results indicate that Ising models constrained with sparsity alone (Ising-L1) yield better fits to functional data than those constrained with SC (Ising-SC). This finding suggests that functional data alone may be sufficient to infer effective brain connectivity for personalized modeling. Given the time and cost associated with dMRI acquisition and processing, this opens the door for more accessible and scalable personalized whole-brain modeling approaches, especially in clinical settings.

The superior performance of Ising-L1 models might be explained by known limitations of dMRI-based tractography. It is well established that dMRI-based tractography underestimates inter-hemispheric, especially homotopic, connections due to limitations in resolving crossing fibers and partial volume effects, and often fails to adequately reconstruct subcortical pathways.^88–90^ Yet, homotopic connections are among the strongest functional links observed in resting-state fMRI, and subcortical structures, such as the thalamus, are essential hubs for coordinating cortical activity and shaping FC.^91, 92^ By relaxing anatomical constraints, Ising-L1 might be able to recover functionally relevant connections directly from BOLD data, which likely contributes to its superior performance at reproducing whole-brain dynamics.

It is important to note that all fMRI data analyzed in this study—real or simulated— were consistently preprocessed using global signal regression (GSR), a widely applied but debated technique in FC studies.^93^ On one hand, GSR helps remove non-neural phenomena such as respiration, cardiac pulsations, scanner drift, and global vascular fluctuations, which are not explicitly implemented in our modeling framework and could otherwise confound FC estimates.^94–98^ Additionally, GSR enhances the detection of anti-correlated networks, which are central to our study.^4, 99^ Furthermore, although it has been stated that GSR may artificially introduce anti-correlations, raising concerns about whether such patterns reflect “genuine” neural dynamics,^100, 101^ several studies have demonstrated that some anti-correlations persist even without GSR, supporting their neurobiological relevance.^4, 99, 102^

Given that the global signal reflects a mixture of neural and non-neural sources outside the scope of our modeling approach, we applied GSR to focus on neural interactions. This is consistent with our modeling limitations, which exclude potential sources of GSR such as cardiac pulsations. However, to assess the robustness of our findings, we repeated the whole-brain simulations using the same fMRI data without GSR (see Supplementary Materials, Section S8), and we showed that the main findings of this study are robust to the use or not of GSR.

While our study provides a novel mechanistic framework for cooperative and competitive interactions in the brain, certain limitations should be acknowledged. First, unlike recent studies that have estimated asymmetric effective connectivity matrices from functional data,^13, 15, 103^ we used symmetric (i.e., bi-directional) long-range connections. The directionality of information flow is fundamental in predictive coding theories, which rely on hierarchical message passing through distinct feedforward and feedback pathways, associated with gamma and alpha oscillations, respectively,^18, 21^ and a proper account of it should improve model performance. Second, we used the same LaNMM in all brain parcels, including subcortical regions. While the LaNMM is designed to capture cortical laminar dynamics, subcortical regions such as the thalamus and basal ganglia exhibit different architectures and dynamics,^92, 104^ and may require specialized models^105^ to fully capture their contributions to whole-brain activity. Future studies should address these limitations to fully integrate the predictive coding principles into the proposed mechanistic model for cooperative and competitive interactions.

## 5 Conclusions

Our findings reinforce that competitive interactions are essential to reproduce key features of large-scale brain activity such as anti-correlations and metastable state transitions. Crucially, we provide a mechanistic substrate for these interactions based on cross-frequency coupling: alpha-power fluctuations suppress gamma-band envelopes, recapitulating invasive laminar recordings in macaque visual cortex and human MEG tasks. Because BOLD signals predominantly reflect metabolic demand tied to gamma-driven synaptic and spiking activity, our envelope-envelope coupling link naturally yields the empirically observed negative correlation between alpha power and BOLD and positive correlation between gamma power and BOLD.

By leveraging maximum-entropy (Ising) models with a simplicity bias to infer effective connectivity independently of the dynamical simulations, we demonstrate that structural priors are not required—and may in fact impede—accurate modeling of whole-brain dynamics. The superior performance of the Ising-L1 models highlights the potential to bypass diffusion MRI, making personalized whole-brain modeling more accessible for research and clinical applications.

Together, the integration of laminar-specific CFC, generative Ising-based connectome inference, and physiologically realistic LaNMM simulations constitutes a novel, mechanistic roadmap for understanding and simulating large-scale brain dynamics, bridging the gap between local oscil-latory circuits and emergent whole-brain activity.

## 6 Acknowledgements

This work has received funding from the European Research Council (ERC) under the European Union’s Horizon 2020 research and innovation programme (Grant Agreement No. 855109; ERC-SyG 2019) and from FET under the European Union’s Horizon 2020 research and innovation programme (Grant Agreement No. 101017716).

Data collection and sharing for the Alzheimer’s Disease Neuroimaging Initiative (ADNI) is funded by the National Institute on Aging (National Institutes of Health Grant U19AG024904). The grantee organization is the Northern California Institute for Research and Education. In the past, ADNI has also received funding from the National Institute of Biomedical Imaging and Bioengineering, the Canadian Institutes of Health Research, and private sector contributions through the Foundation for the National Institutes of Health (FNIH) including generous contributions from the following: AbbVie, Alzheimer’s Association; Alzheimer’s Drug Discovery Foundation; Araclon Biotech; BioClinica, Inc.; Biogen; Bristol-Myers Squibb Company; CereSpir, Inc.; Cogstate; Eisai Inc.; Elan Pharmaceuticals, Inc.; Eli Lilly and Company; EuroImmun; F. Hoffmann-La Roche Ltd and its affiliated company Genentech, Inc.; Fujirebio; GE Healthcare; IXICO Ltd.; Janssen Alzheimer Immunotherapy Research & Development, LLC.; Johnson & Johnson Pharmaceutical Research & Development LLC.; Lumosity; Lundbeck; Merck & Co., Inc.; Meso Scale Diagnostics, LLC.; NeuroRx Research; Neurotrack Technologies; Novartis Pharmaceuticals Corporation; Pfizer Inc.; Piramal Imaging; Servier; Takeda Pharmaceutical Company; and Transition Therapeutics.

## Contributions

Conceptualization, B.M. and GR; methodology, B.M and G.R.; software, all authors; formal analysis, B.M. and M.G.; investigation, B.M., M.G; resources, G.R.; data curation, B.M.; writing—original draft preparation, B.M.; writing—review and editing, B.M and G.R; visualization, B.M.; supervision, G.R.; project administration, G.R.; funding acquisition, G.R., G.K. Data acquisition and curation: ADNI, L.M. and G.K. All authors have read and agreed to the published version of the manuscript.

