## Supplementary Materials for "Bridging local and global dynamics: a biologically grounded model for cooperative and competitive interactions in the brain"

#### S1 LaNMM model equations and parameters.

Neural mass models mathematically represent the dynamics of the average membrane potential and firing rate of various neuronal populations in a cortical column.[1] These models are described by a second-order differential equation that defines the average membrane perturbation $u_{m\leftarrow n}(t)$ a population $m$ experiences due to the synapses from another population $n$ (i.e. the postsynaptic potential). In the synapse-driven formulation,[2] the relationship between the presynaptic firing rate $\varphi_n$ and the postsynaptic potential is governed by the operator $\hat{L}_{m\leftarrow n}$ and its inverse:

$$
\begin{aligned}
u_{m\leftarrow n}(t) &= \hat{L}^{-1}_{m\leftarrow n}\left[C_{m\leftarrow n}\,\varphi_n(t)\right] \\
\hat{L}_{m\leftarrow n}\left[u_{m\leftarrow n}(t)\right] &= C_{m\leftarrow n}\,\varphi_n(t)
\end{aligned}
\tag{1}
$$

Here, $C_{m\leftarrow n}$ is the connectivity strenght between the populations, and $\hat{L}^{-1}_{m\leftarrow n}$ is expressed as a convolution with a kernel $h(t) = Aat\exp[-at]$ for $t > 0$, satisfying the Green's function equation.[3] For generality, synapse $s$ $(m \leftarrow n)$ dynamics are defined as:

$$
\hat{L}_s\left[u_s(t)\right] = \frac{1}{A_s}\left(\frac{1}{a_s}\frac{d^2}{dt^2} + 2\frac{d}{dt} + a_s\right)u_s(t) \tag{2}
$$

where $A_s$ is the synaptic gain, and $a_s = 1/\tau_s$ is the synaptic rate constant. The overall membrane potential perturbation of the post-synaptic neuron, $v_m$, can be calculated as the sum of each

*

pre-synaptic perturbation:

$$v_m(t) = \sum_s u_s(t) \tag{3}$$

and the average firing rate of the neural population, $\varphi_m$, is the output of a sigmoid function:

$$\varphi_m(t) = \frac{2\varphi_0}{1 + e^{r(v_0 - v_m(t))}} \tag{4}$$

Here, $\varphi_0$ is half the maximum firing rate, $v_0$ is the potential at $\varphi_0$, and $r$ determines the sigmoid slope.

Figure S1 shows the neural populations and the synapses between them of the Laminar Neural Mass model. The parameters of the model are described in Table 1. The simulations done with the single and two column models were done using as external inputs Gaussian noise with standard deviation of 5 Hz and mean values of 270 and 90 Hz for $P_1$ and $P_2$ respectively. In contrast, no constant external inputs were used in the whole brain model simulations.

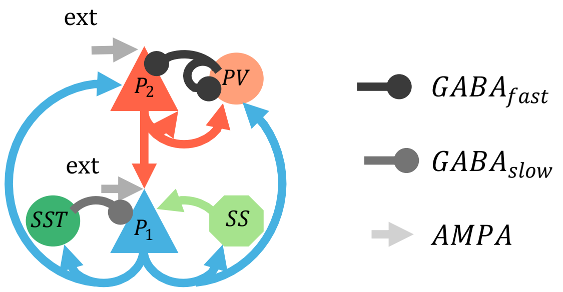


Figure S1: Schematic of the Laminar Neural Mass model showing the neural populations and their connections. The model has two types of AMPAergic synapses depicted with circles and GABAergic synapses depcited with arrows. The model also has two external inputs, one in each pyramidal population.

| **Parameter** | **Description** | **Value** |
|---|---|---|
| $A_s$ | Synaptic gains | $A_{AMPA} = 3.25$ mV<br>$A_{GABA_{slow}} = -22$ mV<br>$A_{GABA_{fast}} = -30$ mV |
| $a_s$ | Synaptic rate constants | $a_{AMPA} = 100\ s^{-1}$<br>$a_{GABA_{slow}} = 50\ s^{-1}$<br>$a_{GABA_{fast}} = 220\ s^{-1}$ |
| $C_{m\leftarrow n}$ | Average number of synaptic contacts from population $m$ to population $n$ | $C_{P_1\leftarrow SS} = 108$<br>$C_{P1\leftarrow SST} = 33.7$<br>$C_{SS\leftarrow P_1} = 135$<br>$C_{SST\leftarrow P_1} = 33.75$<br>$C_{P_2\leftarrow P_2} = 70$<br>$C_{P_2\leftarrow P_1} = 200$<br>$C_{PV\leftarrow P1} = 30$<br>$C_{P_2\leftarrow PV} = 550$<br>$C_{PV\leftarrow PV} = 100$<br>$C_{P1\leftarrow P_2} = 80$<br>$C_{PV\leftarrow P_2} = 200$ |
| $v_0$ | Potential when 50% of the firing rate is achieved | 6 mV* |
| $\varphi_0$ | Half of the maximum firing rate | 2.5 Hz |
| $r$ | Slope of the sigmoid function at $v_0$ | 0.56 $\text{mV}^{-1}$ |

Table 1: Parameters of the Laminar Neural Mass model from Sanchez-Todo et al.[2] used in the study. (*: Except $P_2$, for which $v_0 = 1$ mV)

### S2 Adjustement of the Ising model constraint weights

The frameworks described in the methods section for the generation of generative effective connectivity matrices rely on two constraint weights $\lambda_{SC}$ and $\lambda_{Sp}$. To adjust these constraints, a sweep was done across all subjects in order to assess the goodness of fit of the Ising model as a function of the constraint weights. Te results are shown in Figure S2. In both frameworks, $\lambda$ values were selected to obtain an average PCC between the empirical and the Ising functional connectivity matrices of 0.95. This yielded values of $\lambda_{SC} = 2.5$ and $\lambda_{Sp} = 0.075$. The 0.95 value was selected arbitrarily to ensure that the constraint was enforced without a significant loss in the goodness of fit. The same value was used to fix both constraint weights to allow for a fair comparison between methods.

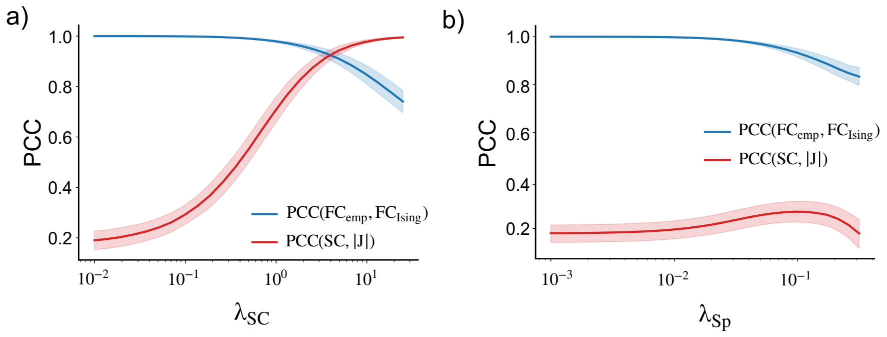


Figure S2: Constraint weight adjustment for the structural connectivity constraint (a) and for the sparsity constraint (b). In blue, Pearson correlation coefficient (PCC) between the empirical functional connectivity and the functional connectivity in the Ising model as a function of the constraint weights $\lambda_i$. Red: PCC between the structural connectome derived from dMRI and Ising $J$ connectivity matrix as a function of the constraint weights $\lambda_i$. The lines show the average value across all subjects and the shaded areas show the standard deviation.

### S3 fMRI-BOLD model

The BOLD time series of each brain region were calculated using the Balloon model proposed by Buxton et al.,[4] according to which the BOLD signal changes depend on two main state variables $q$ and $v$.

$$\frac{\Delta S}{S_0} \cong V_0 \left[ k_1 \left(1 - q(t)\right) + k_2 \left(1 - \frac{q(t)}{v(t)}\right) + k_3 \left(1 - v(t)\right) \right] \tag{5}$$

The state variables $q$ and $v$ represent the deoxyhemoglobin content and the cerebral blood volume respectively. $V_0$ is the resting blood volume fraction in the tissue and $k_i$ are dimensionless coefficients that depend on the MRI acquisition parameters.[5]

The evolution of $q$ and $v$ from the neural activity was calculated using a well-established hemodynamic model.[6] According to this model, in each brain region, the neuronal activity $x(t)$ gives rise to a vasodilatory signal $s$ which is also subject to two feedback regulation terms that depend on itself and on the inflow $f$. The changes over time of the blood inflow are driven by the vaso-dilatory signal $s$ and subsequently cause changes in the blood volume $v$ as well as the deoxyhemoglobin content $q$.

$$\frac{ds}{dt} = x(t) - \kappa s(t) - \gamma(f(t) - 1) \tag{6}$$

$$\frac{df}{dt} = s(t) \tag{7}$$

$$\tau\frac{dv}{dt} = f(t) - v^{1/\alpha} \tag{8}$$

$$\tau\frac{dq}{dt} = f\frac{1 - (1 - E_0)^{1/f}}{E_0} - v^{1/\alpha}\frac{q}{v} \tag{9}$$

where $\kappa$ and $\gamma$ are the rate constants of the signal decay and the feedback regulation by blood flow respectively. The time constant $\tau$ is the mean transit time of blood ( i.e., the average time blood spends in the tissue before leaving through the venous compartments). The magnitude $\alpha$ is the Grubb's exponent that accounts for the stiffness of the venous capillaries.[7] Finally, $E_0$ is the resting state net capillary oxygen extraction rate. The values of all these biophysical parameters can be found in Table S3.

| **Parameter** | **Value** | **Definition, justification and/or source** |
|---|---|---|
| $\gamma$ | 0.41 $s^{-1}$ | Rate constant for auto-regulatory feedback by blood flow[8] |
| $\kappa$ | 0.65 $s^{-1}$ | Rate constant for vaso-dilatory signal decay[8] |
| $\tau$ | 0.98 $s$ | Transit time of blood[8] |
| $E_0$ | 0.4 | Capillary oxygen extraction rate at rest[5] |
| $\alpha$ | 0.32 | Grubb's vessel stiffness exponent[7] |
| $V_0$ | 0.04 | Resting venous blood volume fraction[5] |
| $k_1$ | 2.8 | BOLD coefficient 1 (dimensionless)[5] |
| $k_2$ | 0.57 | BOLD coefficient 2 (dimensionless)[5] |
| $k_3$ | 0.43 | BOLD coefficient 3 (dimensionless)[5] |

Table 2: Parameters used in the BOLD model.

#### S4 Neurovascular coupling models

In the methods section, we introduced a mechanistic model for estimating the metabolic demand of a brain region from the parameters of the generative whole-brain models. This quantity can be used as a proxy for the neural activity that drives the hemodynamic Balloon model presented in the previous section:

$$m(t) = \sum_i \left( \alpha_{GABA} \sum_{j \in exc.} C_{i \leftarrow j} A_{i \leftarrow j} a_{i \leftarrow j} \varphi_j + \alpha_{AMPA} \sum_{j \in inh.} C_{i \leftarrow j} A_{i \leftarrow j} a_{i \leftarrow j} \varphi_j \right) \tag{10}$$

To assess the sensitivity of the BOLD signal to the weighting parameters $\alpha_{GABA}$ and $\alpha_{AMPA}$, we simulated the whole brain dynamics of a representative subject-specific model from our study. Then, for every parcel we calculated the BOLD signals using two different approaches:

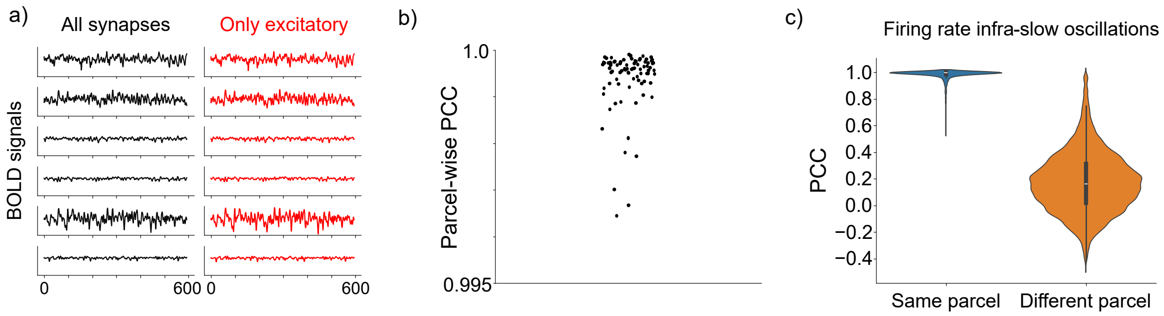


Figure S3: a) Synthetic BOLD traces in various parcels obtained for the same whole brain model using all synapses in the calculation or only the excitatory synapses. b) Pearson Correlation Coefficient (PCC) between the BOLD signals in every parcel of the whole brian model obtained with the two models. c) PCC between the low pass filtered (0.1 Hz cuttoff frequency) firing rates of the neural populations in the whole brain model. In blue the PCC between neural populations of the same parcel, in orange PCC between the neural populations belonging to different parcels.

1) by considering that all synapses contribute equally ($\alpha_{GABA} = \alpha_{AMPA} = 1$) or 2) by considering that only the excitatory synapses contribute to the metabolic demand ($\alpha_{GABA} = 1$ and $\alpha_{AMPA} = 0$).

Figure S3a shows the BOLD signals obtained in a few parcels of the brain for the two approaches. The signals seem identical. In fact, the Pearson correlation coefficients (PCC) between the BOLD signals from the same parcels displayed in Figure S3b show that the difference between both models is minimal (PCC ¿ 0.99). Thus, our results seem robust to the weighting values used for the different synapses of the model.

To understand such low sensitivity to the $\alpha_{GABA}$ and $\alpha_{AMPA}$ parameters, we analyzed the infra-slow oscillations of the different neural populations in the model which are the main driver of the BOLD generation model. Using the same whole brain model, we extracted the firing rates over time of each neural population and we low pass filtered them with a cutoff frequency of 0.1 Hz. Then, we computed pairwise PCC between populations. Figure S3c shows that While the neural populations within the same parcel have highly correlated infra-slow dynamics, neural populations from different parcels show little or no correlation. The activity of the neural populations from the parcel dominates BOLD signal generation due to the larger connectivity strength of the intra-parcel synapses compared to the long-range ones. Thus, regardless of the modeling choice on how to weight them to compute the neural activity, similar BOLD time series are obtained. These findings support the use of a simplified neurovascular coupling model based on excitatory inputs only, as described in the main text.

#### S5 Adjustment of the relative weights of the long range connections

To determine the optimal relative weights of long-range connections (LRCs) for both the cooperative-only and cooperative-competitive modeling approaches, we performed a system-

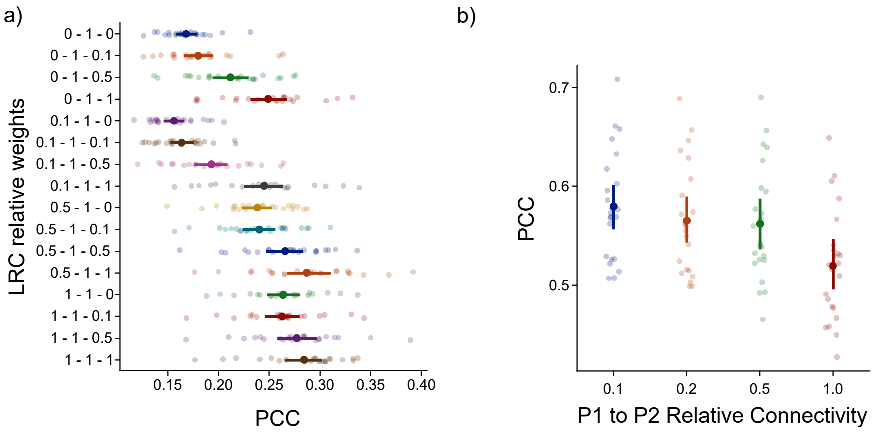


Figure S4: a) Results of the LRC relative weight optimization for the cooperative only models. LRC weights are displayed as $w_{P_1\to P_1} - w_{P_2\to P_2} - w_{P_1\to P_2}$. b) Results of the LRC relative weight optimization for competitive interactions. Each point (translucent) represents an individual subject, and the mean and standard error of the PCC across subjects is also shown (opaque) for each weight configuration.

atic optimization procedure using a subset of 20 subjects randomly selected from the cohort. The optimization aimed to identify the combinations of LRC weights that best reproduce the empirical functional connectivity (FC) patterns, as quantified by the Pearson correlation coefficient (PCC) between empirical and simulated FC matrices as described in the methods section.

For each combination of LRC weights tested, we adjusted the global coupling parameter $G$ individually for each subject, following the same procedure as described in the Methods section. The optimization was performed separately for models based on structural connectomes (cooperative-only models) and Ising-derived effective connectomes constrained by structural connectivity prior, ising-SC (cooperative and competitive models).

In the cooperative-only condition, we considered three types of excitatory-to-excitatory long-range projections: $P_1 \to P_1$, $P_2 \to P_2$, and $P_1 \to P_2$. To explore the space of relative weights systematically, we assessed combinations where the weights could take values in the set $0, 0.1, 0.5, 1$, with the constraint that at least one connection type was set to 1 to avoid redundant combinations. Figure S4a summarizes the results of the optimization procedure for cooperative-only models.

For the cooperative and competitive condition, we focused on the role of competitive interactions specifically in the P1 $\to$ P2 pathway. Here, the cooperative weights for P1 $\to$ P1 and P2 $\to$ P2 were fixed at 1, and we varied the relative weight of competitive P1 $\to$ P2 projections. The results of this optimization procedure are displayed in Figure S4b.

#### S6 Spatiotemporal analysis for various cluster numbers

To assess the robustness of the spatiotemporal dynamics analysis based on the Leading Eigenvector Dynamics Analysis (LEiDA), we evaluated how the Kullback–Leibler (KL) divergence and entropy rate (ER) differences between empirical and synthetic data vary when the number

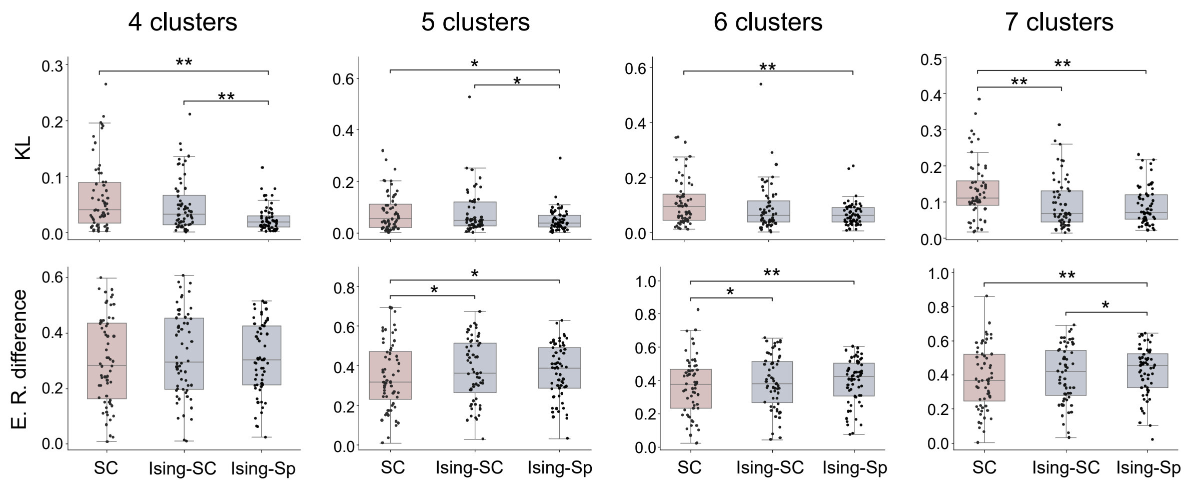


Figure S5: Kullback–Leibler distance (KL) and Entropy rate difference between synthetic and empirical data for the different groups and number of clusters. (*: $p < 0.05$, **: $p < 0.01$ with Wilcoxon signed-rank test)

of functional connectivity (FC) states (clusters) is changed.

While the main analysis focused on the optimal clustering solution with 3 FC states, identified through multiple clustering quality metrics and correspondence with canonical functional networks (see Methods and Results), here we extend the analysis to cluster numbers ranging from 3 to 7 to examine whether our findings generalize beyond the optimal solution.

Figure S5 summarizes the results. The statistically significant differences between the KL of the SC and ising-Sp groups is robust to the number of clusters used in the analysis. However, the difference beteween the ising-SC and the ising-Sp groups is attenuated for larger cluster numbers. In contrast, the SC and Ising-SC groups only display statstically significant differences for 7 clusters.

Regarding entropy rate differences, we found that all modeling approaches showed comparable performance when using 3 and 4 clusters, with no statistically significant differences across groups. However, when the number of clusters is increased the SC group shows significantly lower values compared to the other groups.

#### S7 Stochastic and Deterministic whole brain models

While stochastic noise is commonly added as input to neural mass models, throughout this study we used deterministic generative whole brain models (i.e. without any stochastic drive). To assess the impact of this choice we produced deterministic and stochastic subject-specific whole-brain models for all subjects using the Ising-L1 connectomes. To do so, for each subject we adjusted the Global coupling using either constant values or Gaussian noise with a standard deviation of 5 Hz as external inputs to the $P_1$ and $P_2$ populations in each parcel.

Similar PCC values were obtained for the best fits using the deterministic and stochastic models as shown in Figure S6a. In addition, Figure S6b shows that the FC matrices of both models

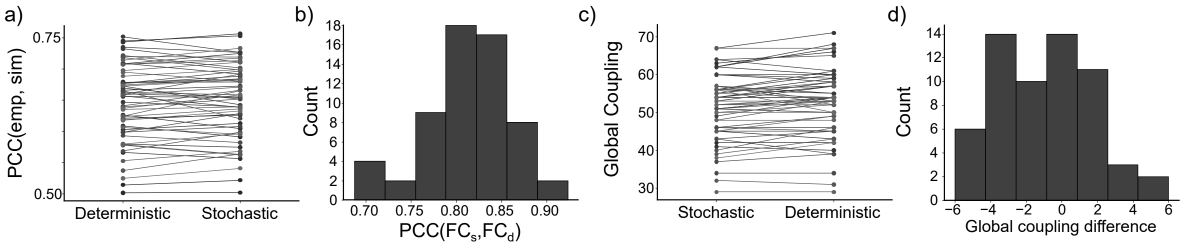


Figure S6: a)Pearson's correlation coefficient (PCC) between the empirical and the synthetic FC matrices for all subjects using deterministic and stochastic models. b) PCC between the FC matrices of the stochastic and deterministic models for all subjects. c) Global coupling values of the best fit to the empirical data for the stochastic and deterministic models. d) Global coupling difference between the stochastic and deterministic models for all subjects.

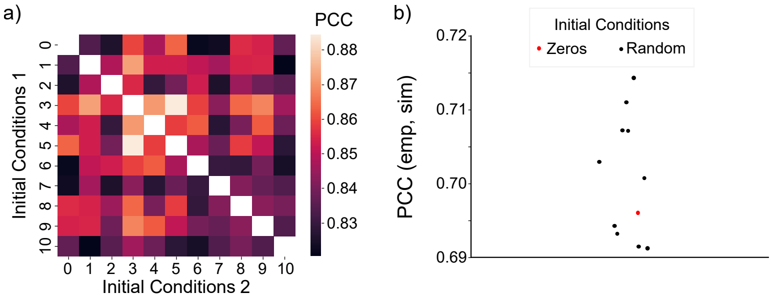


Figure S7: a) Pearson's correlation coefficient between the FC matrices obtained using different initial conditions. Index 0 corresponds to the case with all variables set to 0 and the rest correspond to the variables initialized using random values. b) PCC between the empirical and the

display a high correlation among subjects (mean PCC of 0.81). In fact, the Global coupling values that best fit the empirical data are similar in both models as shown in Figure S6c and Figure S6d. Based on these results we can conclude that similar fits to the empirical data can be obtained regardless of the use or not of stochastic noise in the whole-brain models. In addition, the model parameters that best fit the empirical data are very similar in both cases. Thus, the results are robust to the use or not of stochastic noise inputs.

The results of the deterministic models may vary depending on the initial conditions chosen to run the simulations. Throughout this study, all models were arbitrarily initialized with all the internal variables set to 0. To assess the impact of this choice, we simulated the FC matrix using a representative subject-specific model from our study with 10 different initial conditions. Figure S7a shows the pairwise PCC between the FC matrices obtained in the different simulations and Figure S7b shows the PCC between these simulations and the empirical FC of the subject. The different simulations display a high correlation (average PCC of 0.85) and their correlations with the empirical Fc have similar values. Thus, while the results depend on the initial conditions, overall this does not seem to impact the conclusions of the study.

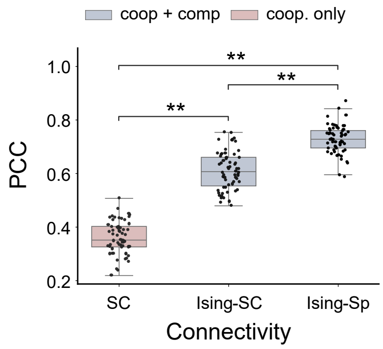


Figure S8: Pearson's correlation coefficient between the empirical and the synthetic FC matrices for the three types of connectivity matrices. (**: $p < 0.01$ with Wilcoxon signed-rank test)

### S8 Subject specific generative models without GSR

To assess the robustness of the results to the usage or not of GSR in the fMRI pre-processing, we created generative whole brain models without GSR for the same cohort of subjects. The workflow to create the subject-specific whole brain models was exactly the same. The only difference was that GSR was not used in the pre-processing of the fMRI data. The overall fitting results for the three families of connectivity matrices are displayed in Figure S8
